# L-DOPA Reduces Model-Free Control of Behavior by Attenuating the Transfer of Value to Action

**DOI:** 10.1101/086116

**Authors:** Nils B. Kroemer, Ying Lee, Shakoor Pooseh, Ben Eppinger, Thomas Goschke, Michael N. Smolka

## Abstract

Dopamine is a key neurotransmitter in reinforcement learning and action control. Recent findings suggest that these components are inherently entangled. Here, we tested if increases in dopamine tone by administration of L-DOPA upregulate deliberative “model-based” control of behavior or reflexive “model-free” control as predicted by dual-control reinforcement-learning models. Alternatively, L-DOPA may impair learning as suggested by “value” or “thrift” theories of dopamine. To this end, we employed a two-stage Markov decision-task to investigate the effect of L-DOPA (randomized cross-over) on behavioral control while brain activation was measured using fMRI. L-DOPA led to attenuated model-free control of behavior as indicated by the reduced impact of reward on choice and increased stochasticity of model-free choices. Correspondingly, in the brain, L-DOPA decreased the effect of reward while prediction-error signals were unaffected. Taken together, our results suggest that L-DOPA reduces model-free control of behavior by attenuating the transfer of value to action.

## Introduction

Goal-directed actions require considerations along at least two axes. First, action policies reflect which sequence of actions ultimately leads to reinforcement, thereby guiding the allocation of response vigor ^1^. Second, across repeated decisions, estimates of the reward rate have to be updated to learn about potential changes in the environment. Whereas the associated computations recruit partly divergent brain networks, the information is integrated in a common region: the nucleus accumbens (NAcc). The NAcc acts as a limbic–motor interface and has been referred to as “nexus of goals” ^4^ While there is mounting evidence each supporting the role of mesocorticolimbic dopamine in the control of vigor^5-7^ and reinforcement learning^8, 9^, little is known about how these two aspects go hand in hand in order to support dynamic control of behavior. Hence, we sought to test how increases in dopamine tone via the administration of L-DOPA affect the correspondence between reinforcement learning and goal-directed behavior.

An important distinction between the action-control modes of habitual vs. goal-directed behavior has been made in dual-action choice models of reinforcement learning^10^. On the one hand, “model-free” (MF) control appears as reflexive because it is based on direct reinforcement of successful actions via reward prediction errors (RPE), putatively encoded in mesocorticolimbic dopamine neurons^11, 12^. On the other hand, “model-based” (MB) control appears as deliberative because it is based on a learned model of the task structure, which allows agents to plan actions ahead of time by simulating their potential outcome^3, 13, 14^ Since MB control operates on “goal values”, it has been associated with goal-directed behavior. In contrast, MF control operates on “habit values”, which are divorced from the goal of reinforcement^15^.

Recent work in rodents has demonstrated that action control and reinforcement learning are indeed entangled in dopamine signaling and that RPEs are contingent on action^16,17^. Phasic dopamine at the beginning of a trial invigorates actions whereas it reinforces actions at the end of a trial when the outcome is being presented^16, 18^. Thereby, dopaminergic RPEs act as “teaching signals”^18^. The amplitude of phasic RPE signals is dependent on dopamine as increasing (via L-DOPA) versus reducing (via haloperidol) dopamine transmission leads to corresponding changes in RPE BOLD signals in humans^19^ Furthermore, Guitart-Masip et al.^20^ have shown that the mesocorticolimbic system tracks instrumental values in opposite ways for go vs. no-go decisions, indicating that value 17 representations are gated by action in accordance with animal research^17^. In addition to phasic signaling, dopamine tone has been hypothesized to track the average reward rate^1^ and plays an important role in facilitating response vigor^21, 22^. Dopamine has also been associated with the arbitration between action-control modes^13, 14^, but the differential contributions of tonic versus phasic signaling to arbitration are not well understood in humans. Collectively, this evidence of essential crosstalk between action control and reinforcement learning raises the fundamental question how dopamine signals integrate and balance both aspects adaptively.

Depending on which theory one ascribes to when formulating behavioral predictions, one might arrive at conflicting expectations of how exogenous dopamine modulation would impact action and learning^23^. As exogenous modulation of dopamine, we used the synthetic precursor L-DOPA, which increases postsynaptic levels of dopamine and leads to an increase in tone^24^. In a behavioral study, Wunderlich, Smittenaar and Dolan^13^ have shown that L-DOPA leads to an upregulation of deliberative MB control without altering learning rates. However, according to reinforcement learning theories, L-DOPA would be expected to increase RPE amplitudes^19^, at least for positive outcomes (because high dopamine tone may impair negative outcome learning via D2 receptors^25, 26^). Alternatively, L-DOPA may impair learning, which is suggested by both “value”^16^ and “thrift” theories^5^ of dopamine function. The value theory^16^ proposes that heightened dopamine tone would reduce the local change in dopamine, thereby reducing the impact of phasic RPEs on action control^23, 27^. Moreover, according to the thrift theory^5^, increased dopamine tone reduces the degree to which learned reward values affect behavioral choice, making actions effectively more independent of (learned) value. Consequently, both value and thrift theories make highly similar behavioral predictions although they emphasize complementary aspects of dopamine function. Since a behavioral study cannot conclusively address if changes in MB control arise due to changes in RPE signals in the mesocorticolimbic system or, alternatively, a difference in the correspondence between RPEs and behavior, we sought to bridge this essential gap in our current study.

Therefore, in line with the suggestion by Collins and Frank^23^, we tested common theories of dopamine function using an exogenous increase in tonic dopamine while recording endogenous responses to rewarding events. Critically, we used a two-stage Markov task that has been designed to distinguish MF and MB control, including their RPE correlates, according to the dual-action choice framework of reinforcement learning (Figure 1a, see 3 10 methods)^3, 10^. This enables us to separate reflexive from prospective facets of reinforcement learning and actions control that may be weighed differentially after administration of L-DOPA^13^. Furthermore, we can assess if effects of L-DOPA provide support for the reinforcement learning (i.e., an improvement via facilitation of RPE signaling) or the value/thrift account of dopamine function (i.e., a reduced correspondence between learned values and action). Hence, we focused the analyses a priori on regions in the mesocorticolimbic system where MF and MB RPE signals are commonly observed (for details, see methods).

**Figure 1:**
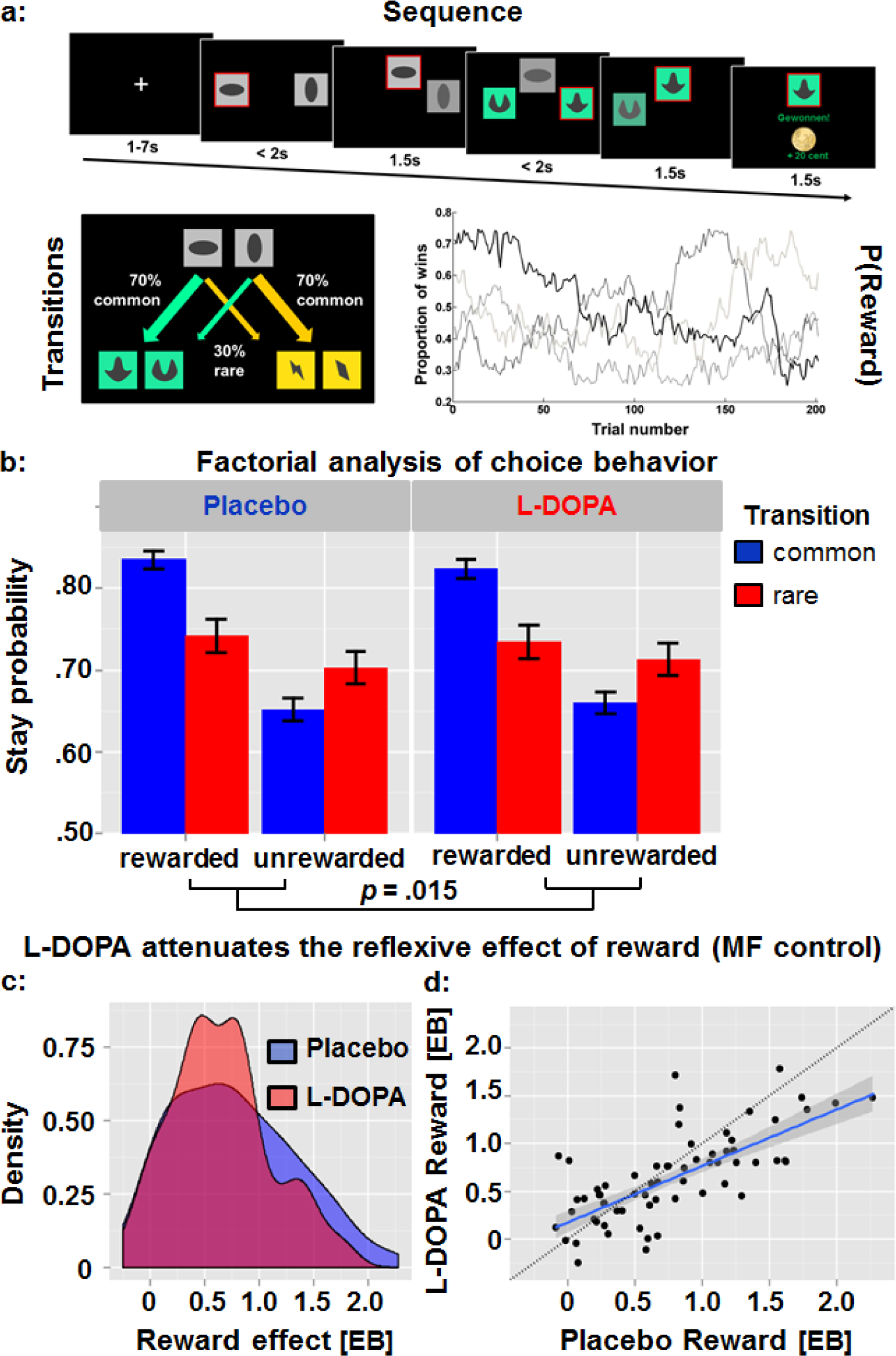
A: Schematic of the task sequence in one trial, transition contingencies, and random walks of the probabilities to win for the four options (green/yellow boxes at stage 2). B: The factorial analysis of stay/switch behavior indicated that L-DOPA attenuated the effect of preceding reward on stay probability of the next trial, which is a signature of model-free (MF) control of behavior. Error bars depict 95% confidence intervals at the trial level. C: Density plots of the distribution of empirical Bayes (EB) parameter estimates of the preceding reward effect on stay/switch behavior. Inspection of the density and scatter plots (d) indicates that L-DOPA “shrinks” strong model-free control of behavior during placebo towards zero, leading to an overall decrease in the impact of reward on stay probabilities. Blue line = robust linear fit, black line = identity.

## Results

### L-DOPA reduces the effect of reward on stay/switch behavior

In line with previous studies^3, 13^, we estimated the main effects of reward, transition, and the Reward x Transition (RxT) interaction on the repetition of the same choice at the first stage of the next trial (“stay”) using full-mixed effects modeling of placebo and drug sessions (see methods). The obtained individual estimates of the effects were than compared between conditions and used for further correlational analyses. Within this hierarchical logistic regression analysis, the main effect of reward corresponds to MF control (i.e., the reflexive effect of reward regardless of transition) whereas the RxT interaction term captures the use of the transition structure in the goal-directed pursuit of reward, corresponding to MB control.

Administration of L-DOPA attenuated the effect of preceding reward on stay probabilities of the next trial (*t* = -2.50; *p* = .015; Fig. 1b-d) thereby reducing the reflexive MF facilitation of response after reward. An equivalent yet less powerful non-parametric test (i.e., Wilcoxon signed rank test on OLS estimates disregarding group priors) yielded a slightly attenuated estimate of the drug effect on MF control, which was only significant at trend level (*Z* = -1.74; Monte Carlo *p* = .084 95% CI [.079-.089]). Descriptively, receiving a reward increased the odds of repeating the same choice at stage 1 of the next trial by 116.8% during placebo sessions (*b*_*P.REW*_ = .774) whereas it only increased the odds by 91.8% during L-DOPA sessions (*b*_*D.REW*_ = .651). In contrast, L-DOPA had no significant effect on MB control as indicated by RxT (*p* = .27) or the intercept, which reflects a general bias to repeat the previous choice (at centered reward and transition terms, *p* = .75).

### L-DOPA reduces model-free control at second stage in the computational model

To provide a more nuanced assessment of drug effects on reinforcement, we set up the seven parameter computational model (M_7P_) proposed Daw, Gershman, Seymour, Dayan and Dolan^3^ and used maximum a posteriori (MAP) as fitting algorithm to better approximate the normal distribution of model parameters for parametric statistics^14^. L-DOPA increased stochasticity of choices at the second stage (i.e., reduced *β2*), which is primarily under MF control (*p* = .008). This significant difference was also seen in the non-parametric Wilcoxon signed rank test (*Z* = -2.80; Monte Carlo *p* = .004 95% CI [.003-.005]). Notably, L-DOPA appeared to have opposite effects on stochasticity, leading to a positive Stage x Drug interaction (*p* < .001; Figure 2a) despite the positive correlation between *β*s at both stages (*r*_*D*_ = .43, *p* < .001; *r*_*P*_ = .36, *p* = .004).

**Figure 2:**
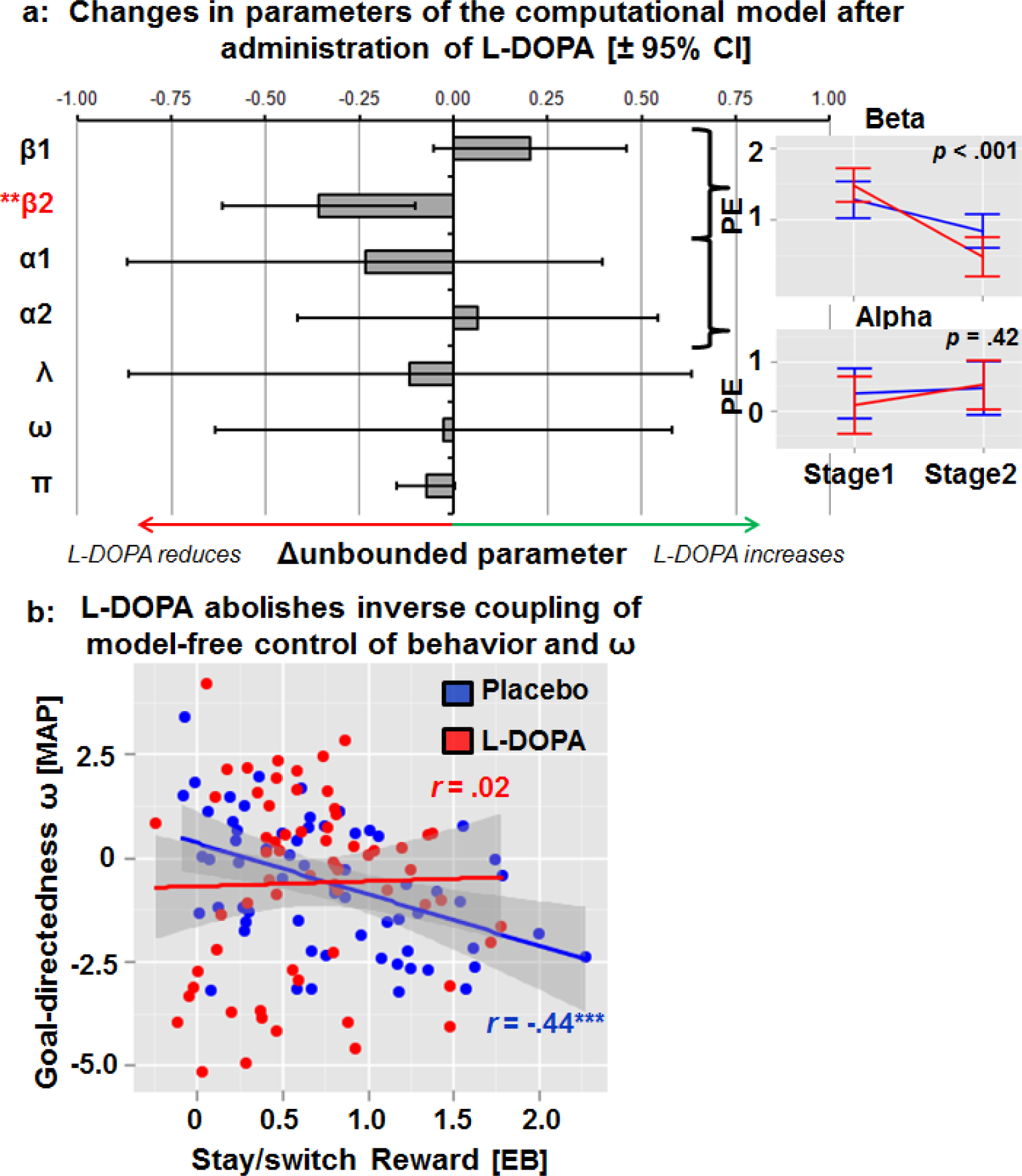
Administration of L-DOPA increased stochasticity of choices at the second (i.e., primarily model-free) stage and abolished the negative correlation between goal-directedness in behavior, ω, with model-free control (i.e., the preceding reward bias on stay/switch behavior). A: L-DOPA reduces *β2*, which reflects the consistency of choices with values at stage 2 (*t* = -2.737; *p* = .008). Interactions of L-DOPA with task stage on *α* and *β* parameters are shown in the inset graph. B: Whereas there is a strong negative correlation between model-free control of behavior and goal-directedness during placebo visits, the correlation is abolished by L-DOPA, indicating that the interaction between model-free and model-based control systems was altered by increases in dopamine tone (*p* = .046). EB = empirical Bayes, PE = parameter estimate

In line with the logistic regression analysis of stay probabilities, L-DOPA did not affect the degree of MB control, which is captured by the weighting parameter ω in the computational model (*p* = .94). Instead, L-DOPA abolished the negative correlation between MB control weighting parameter ω and the individually estimated MF control by preceding reward (*r*_*D*_ = .02, *p* = .86; *r*_*P*_ = -.44, *p* < .001; significant drug interaction term in HLM, *p* = .046; Figure 2b) making both action control modes operate more independently. No main or interaction effect of drug was seen for learning rates at both stages and all other parameters were not significantly different, although in line with previous research^13, 28^, L-DOPA tended to increase switching as captured by the repetition bias parameter (*p* = .067; Figure 2). Lastly, there were no significant effects of gender or order in factorial or computational analyses of behavior.

Due to the fact that we could not replicate the increase of MB control after administration of L-DOPA reported in Wunderlich, Smittenaar and Dolan^13^, we tested if drug effects were dependent on overall model fit. First, L-DOPA did not significantly alter the log likelihood (LL) of the computational model (*t* = 1.50, *p* = .15). Second, we added LL_M7_._D_ to repeated measures ANOVAs where estimates of MB control (RxT or ω) were the dependent variables, respectively. Administration of L-DOPA increased the RxT interaction term when the model fit was high, *F*(1,63) = 10.92, *p* = .002. A similar trend was observed for ω, *F*(1,63) = 2.84, *p* = .097. In other words, increases in MB control elicited by L-DOPA might be dependent on how well the model captures behavior. To test if the differences in drug effects might be driven by differences in the characteristics of the selected samples (i.e., undergraduates vs. a representative of adults), we used a working memory score computed from the operation span task (for details, see SI). Higher working memory capacity was associated with better average model fit (*p* = .015) and increases in MB control after L-DOPA (ω: *p* = .003; RxT: *p* = .089). These results indicate that the effects of L-DOPA are partly dependent on cognitive abilities such as working memory capacity. Critically, reduced MF control of behavior after administration of L-DOPA as evidenced by the effect of preceding reward or *β2* was found independently of model fit and working memory capacity.

Furthermore, we examined alternative models tested by Wunderlich, Smittenaar and Dolan^13^. However, the models were inferior in terms of overall model fit and we found no consistent effect of L-DOPA on MB behavior or reward learning (see Table S.1, SI). Lastly, similar effects of L-DOPA on *β2* (*p* = .013; Wilcoxon signed ranks test *Z* = -2.75; Monte Carlo *p* = .006 95% CI [.004-.007]) were also obtained with a recent reparametrization of M_7P_ where ω is replaced by separate *β*s for MF and MB control^29-31^. indicating that the obtained result is robust to minor changes in the setup of the computational model.

To summarize, L-DOPA increases stochasticity of choices at the second stage of the task which is primarily under MF control. In contrast, drug effects on MB control were inconsistently found across models and not evident in the best fitting model. Furthermore, potential increases in MB control following the administration of L-DOPA were dependent on high overall model fit or working memory capacity. Taken together, the results of the computational models echo the results of the factorial analysis and point conclusively to reduced MF control, but largely unaffected MB control (on average) after administration of L-DOPA.

### No effect of L-DOPA on BOLD correlates of RPE signals

To test for effects of L-DOPA on BOLD response correlates of RPE signals, we set up two separate second-level statistics for MF RPE and MB RPE contrast images including both conditions as repeated measures factor while controlling for order as a covariate. In line with previous studies^3^, we observed a widely distributed brain network tracking MF RPE signals encompassing the ventral and dorsal striatum, the dopaminergic midbrain, the orbitofrontal cortex, the dorsolateral prefrontal cortex, and the posterior cingulate cortex (Figure 3; Table S.2). Contrary to MF RPE, MB RPE signals were found to be spatially sparser encompassing the ventral striatum (where overlap with MF RPE signals occurs) and vmPFC only. These results replicate the results reported by Daw, Gershman, Seymour, Dayan and Dolan^3^.

**Figure 3:**
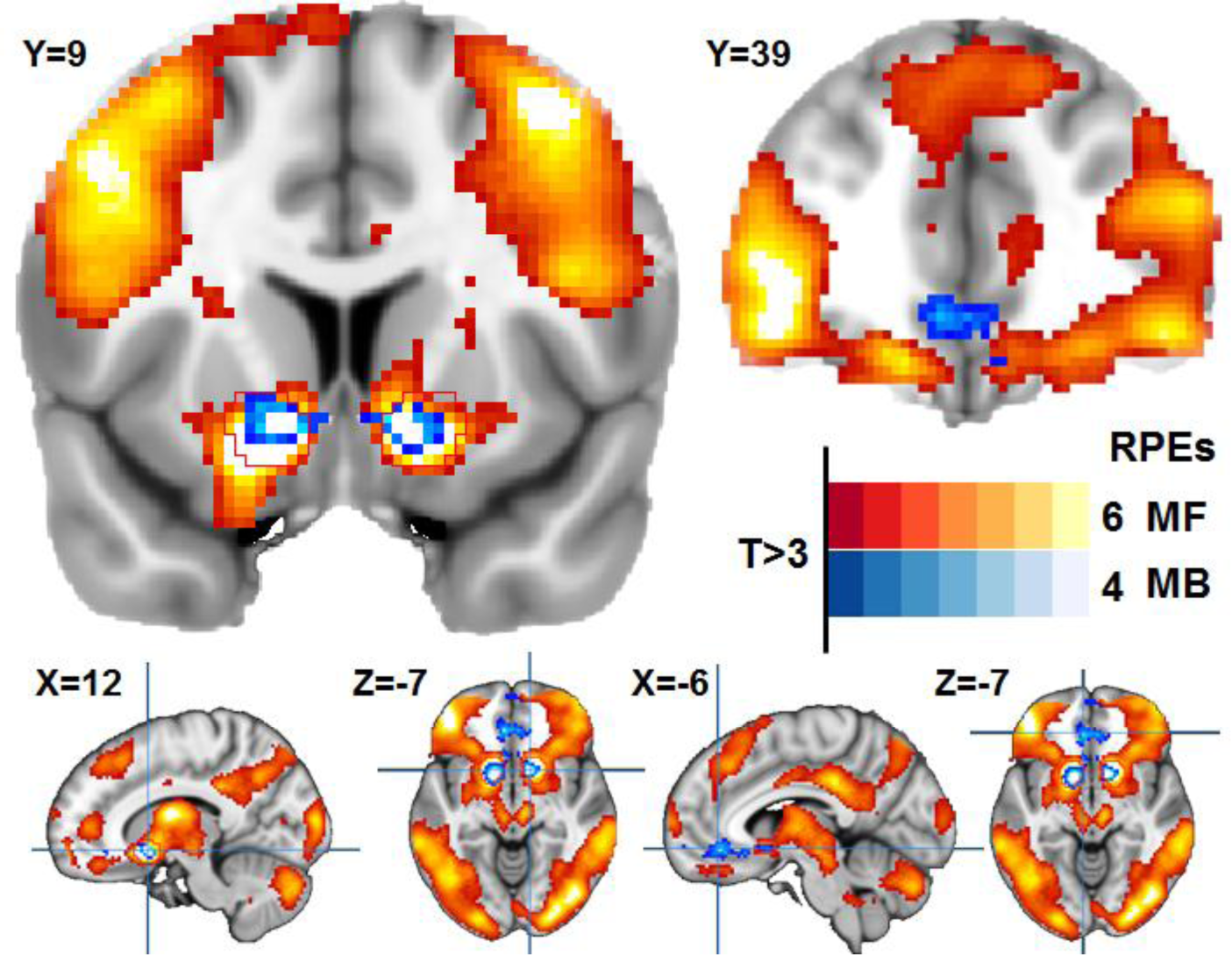
Model-free (MF; red-yellow) and model-based (MB) reward prediction error signals (RPE; blue-white) pooled for placebo and L-DOPA sessions (with session order as covariate). In line with Daw et al. (2011), overlap of MF and MB reward prediction error signals were found in the nucleus accumbens (ROI outline shown in red) and ventromedial prefrontal cortex.

In contrast to predictions of the reinforcement learning theory, yet in line with the absence of behavioral effects on learning rates, we observed no modulatory effects of L-DOPA on RPE signals (Figure 4). At a whole-brain level, no differences survived a correction for multiple comparisons (cluster-forming threshold *p* < .001) and drug effects were virtually absent even at an uncorrected cluster extent threshold (see SI). Likewise, at the level of a priori ROIs, we observed no effects of L-DOPA (*ps* > .1). This absence of a drug effect was further corroborated by an additional time course analysis: We concatenated sessions and participants to improve the estimation of potential drug effects across the group, similar to the behavior analysis, but failed to see a modulatory effect of L-DOPA on RPE signaling (|*t*| < 1.21, *p* > .23). Whereas there was no effect of L-DOPA on average parametric effects, we found an increase in interindividual variability of the MF RPE signals in the NAcc during the L-DOPA session, F(64,64) = 1.61, *p* = .030, which would, however, not survive correction for multiple comparisons across ROIs and/or contrasts. To summarize, across whole-brain and a priori ROIs analyses, there was no indication of an altered correspondence between BOLD response and RPE signals after administration of L-DOPA.

**Figure 4:**
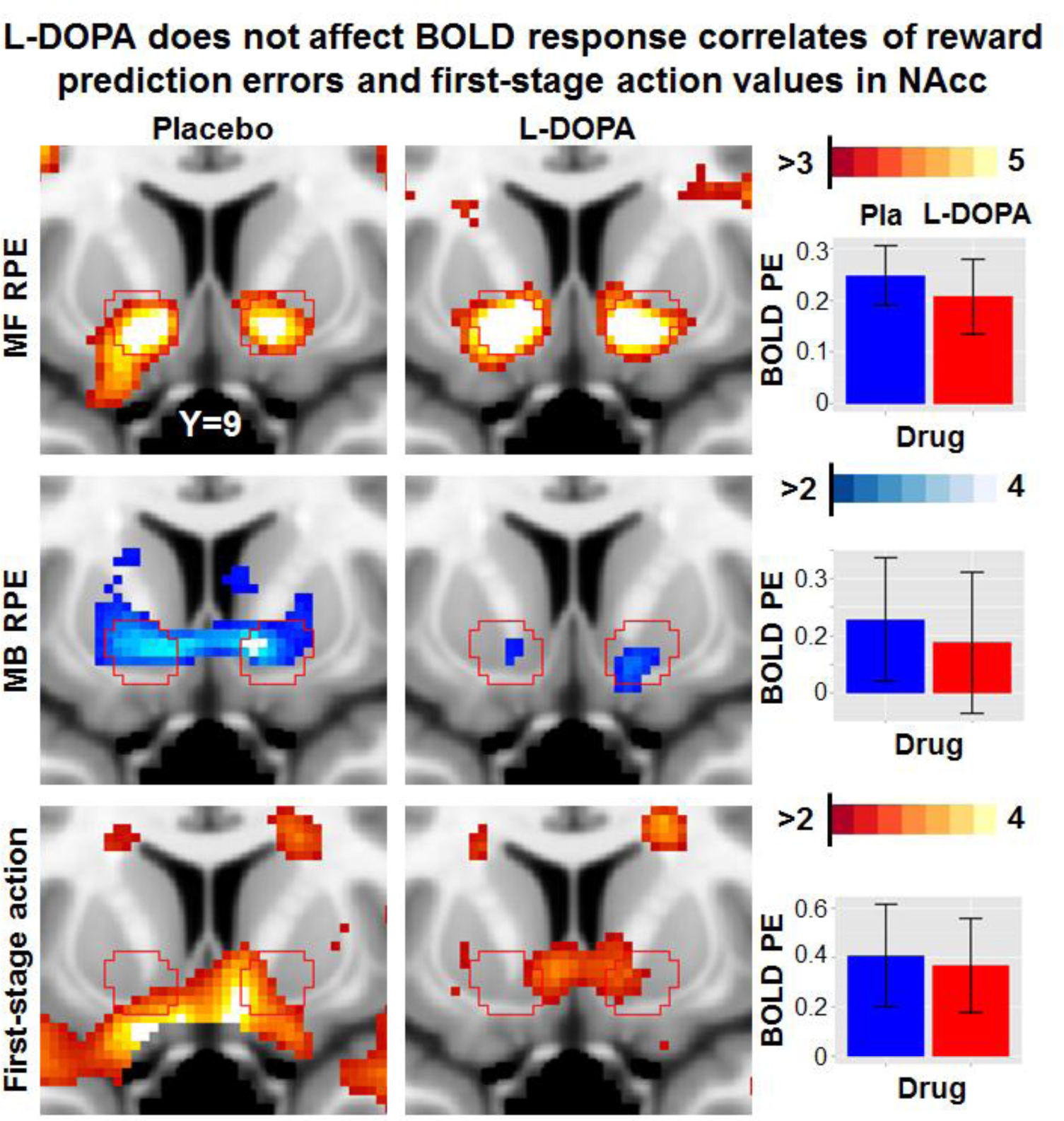
A priori region-of-interest (ROI) sections of the nucleus accumbens (NAcc) for the parametric effects of interest model-free (MF) and model-based (MB) reward prediction error (RPE) signals and first-stage action values depicted separately for placebo and drug sessions. Inset graphs show extracted parameter estimates (PE) and 95% confidence intervals (CI). Likewise, non-significant differences were obtained in ROI analyses for the ventromedial prefrontal cortex and in exploratory whole-brain analyses. Pla = placebo, PE = Parameter estimate

### L-DOPA reduces the effect of reward at the outcome and subsequent trial-onset stage

In the factorial analysis and the computational model of behavior, we found that L-DOPA reduced reflexive MF control by reward without altering the encoding of RPEs. Hence, we set up an alternative first-level model of the task to assess the simple main effect of rewarded vs. unrewarded events. Second-level statistics were computed separately for the outcome stage and the subsequent trial onset, where participants may decide to stay or switch depending on the outcome and the transition of the previous trial. As a covariate, we included the individually estimated effect of reward obtained in the factorial analysis of stay/switch behavior conducted at this stage of the task (Figure 1c-d). This parameter captures the degree to which individuals reflexively adjust their first-stage choice after receiving a reward. Based on the behavioral results, which were in line with the predictions of the thrift and value theories of dopamine, we hypothesized that L-DOPA would reduce the effect of reward (rewarded – unrewarded events) on BOLD response in the mesocorticolimbic ROIs VTA/SN, NAcc, and vmPFC.

Across the three ROIs, we found a significant effect of reward (*p* < .001), no significant effect of drug (*p* = .23), but a significant interaction Drug x Reward (*p* = .027; Table S.3). This interaction effect was driven by a reduced contrast between rewarded and unrewarded events during L-DOPA vs. placebo visits (Figure 5a; Figure S.2). Additional single ROI models indicated that the Drug x Reward effect was significant within the VTA/SN (*p* = .014; survives correction across 3 ROIs) and vmPFC (*p* = .047), but not in the NAcc alone (*p* = .32). Analogous effects were obtained with a voxel-based approach and coordinates of peak effects are reported in the SI. No other effects were seen in a whole-brain analysis that survived correction for multiple comparisons and there was no general effect of reward at the outcome stage on grey matter BOLD response (*t* = -.482, *p* = .63).

**Figure 5:**
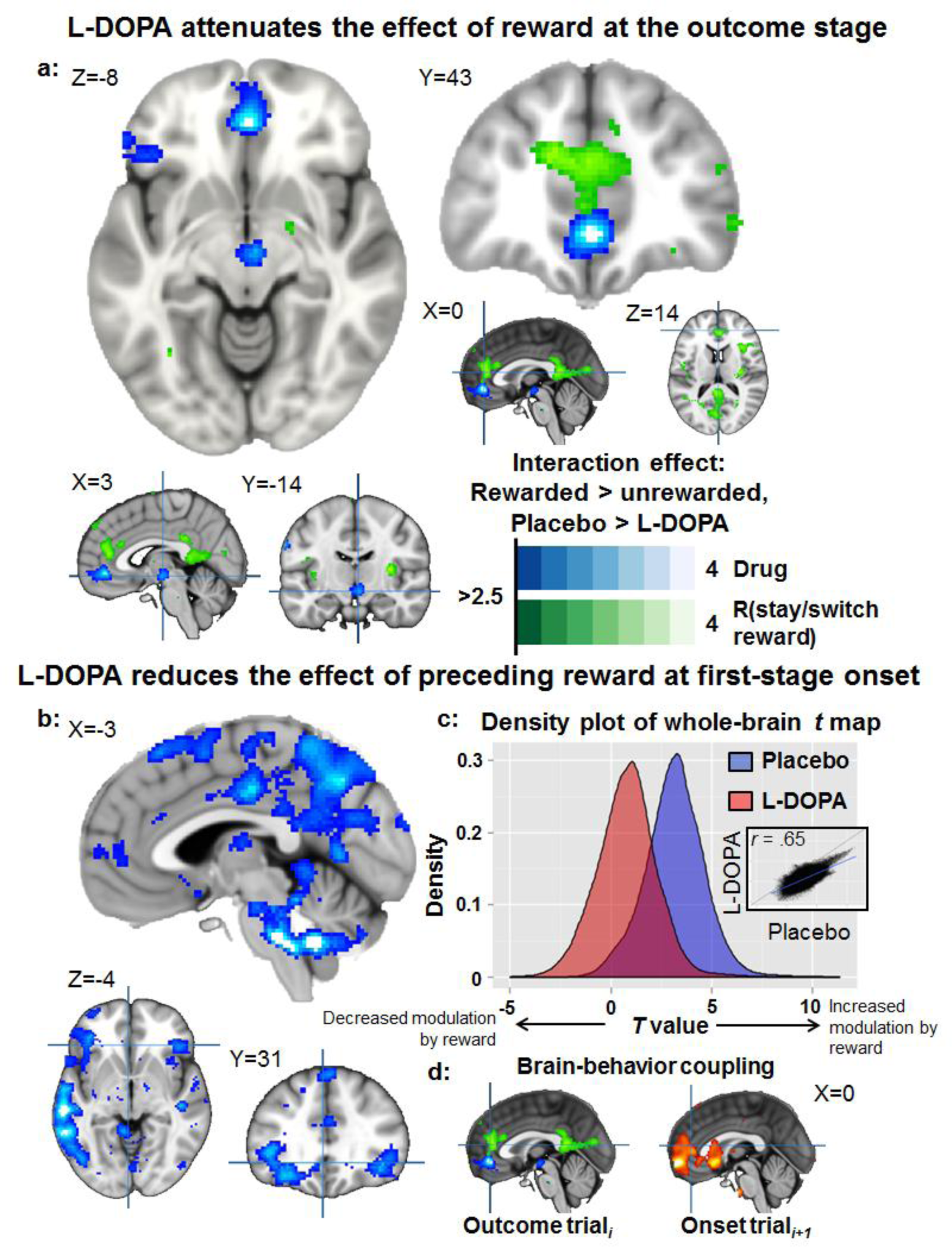
A: L-DOPA reduced the effect of reward on brain activation in the ventral tegmental area and the ventromedial prefrontal cortex (blue color map) at the outcome stage. Moreover, it reduces the correlation of brain activation with the reflexive effect of reward on behavior (derived from stay/switch analyses) in an adjacent cluster of the anterior cingulate cortex (drug interaction within the ROI *p* = .004). This indicates that the transfer of model-free values to action is reduced. B: In line with the attenuating effect at the outcome stage, L-DOPA reduces the effect of preceding reward on brain activation at the onset of the next trial throughout the brain (c). The inset scatterplot depicts that the group t-values are reproducible across conditions at a voxel-by-voxel level, but that the preceding reward effect is globally attenuated (grey matter *p* = .003). D: At first-stage onset, stronger activation in the prefrontal cortex is associated with the effect of reward on behavior regardless of drug condition (hot color map; scaled analogous to the other contrasts and plotted next to the image of the Drug x Reward Bias interaction in a).

Likewise, at the onset of the following trial, we observed a significant decrease of the reward effect on BOLD response across the three a priori ROIs after administration of L-DOPA (*p* = .019). At this stage, however, the effect was not only restricted to the mesocorticolimbic system because we found a significant modulation at the whole-brain level across grey matter voxels (*t* = -3.034, *p* = .003; Figure 5b).

Lastly, to evaluate if L-DOPA affects the coupling between behavior and reward signals, we assessed if L-DOPA reduced the correspondence between MF control as captured by the reflexive effect of reward and BOLD response to reward. Here, we observed a reduced correlation between the neural and the behavioral reward effect in the ACC (*r*_*P*_ = .36, *p* = .003; *r*_*D*_ = -.18, *p* = .15; drug interaction term in HLM *p* = .004; voxel-based *t*_*max*_ = 3.38, *p_SVC_* = .056; 2/38/12), which would survive correction for multiple comparison across ROIs and stages (α < .0083). At the first stage of the following trial, we found that the neural effect of preceding reward corresponded with the reflexive effect of reward in the vmPFC regardless of the drug condition (*r_P_* = .24, *p* = .053; *r*_*D*_ = .34, *p* = .006, no significant drug interaction; *t*_*max*_ = 4.60, *p*_*SVC*_ = .002; *k* = 605, *p*_*FWE.clust*_ = .002; -4/60/-6; extending to the ACC ROI *t*_*max*_ = 3.52, *p*_*SVC*_ = .035; -2/48/6). Furthermore, we observed a significant correlation in the orbitofrontal gyrus outside of our a priori ROIs (*t*_*max*_ = 4.64, *k* = 511, *p*_*FWE.clust*_ = .004; 20/32/-8).

To summarize, L-DOPA reduced the correspondence between the neural reward effect at the outcome stage and the behavioral reward effect in the ACC. Moreover, when participants were facing the next choice, the facilitating effect of reward on BOLD response was reduced throughout the brain, but the correspondence between the neural reward effect and behavior was not affected by L-DOPA. This suggests an attenuated net effect of the reward. Hence, this corroborates the observed decrease in the reflexive effect of reward on behavior.

### L-DOPA facilitates exploratory behavior

Whereas both value and thrift theories predict that increases in dopamine tone reduce MF control as we have observed, one of the key predictions of the thrift theory is that heightened dopamine tone will facilitate exploratory behavior. To test this hypothesis, we used multilevel analyses of RT as well as first- and second-stage action values. We hypothesized that exploration would be indicated by less deterministic choices (“noisy” exploration^32^) and higher RT for switches (“strategic” exploration^32^).

In line with the strategic aspect of exploration, we observed that L-DOPA increased the discrepancy in RT between stay and switch trials at stage 1 (Wilcoxon signed ranks test *Z* = -2.565, Monte Carlo *p* = .011 95% CI [.009, .013]), which was mainly driven by slower RT during switch trials in L-DOPA sessions (RT_D.switch_ = 0.781 s vs. RT_P.switch_ = 0.765 s; RT_D.stay_ = .739 s vs. RT_P.stay_ = .735 s). Notably, there was no general effect of L-DOPA on RT at first stage and no shift in the distribution of the probability of choices (Figure 6a).

**Figure 6:**
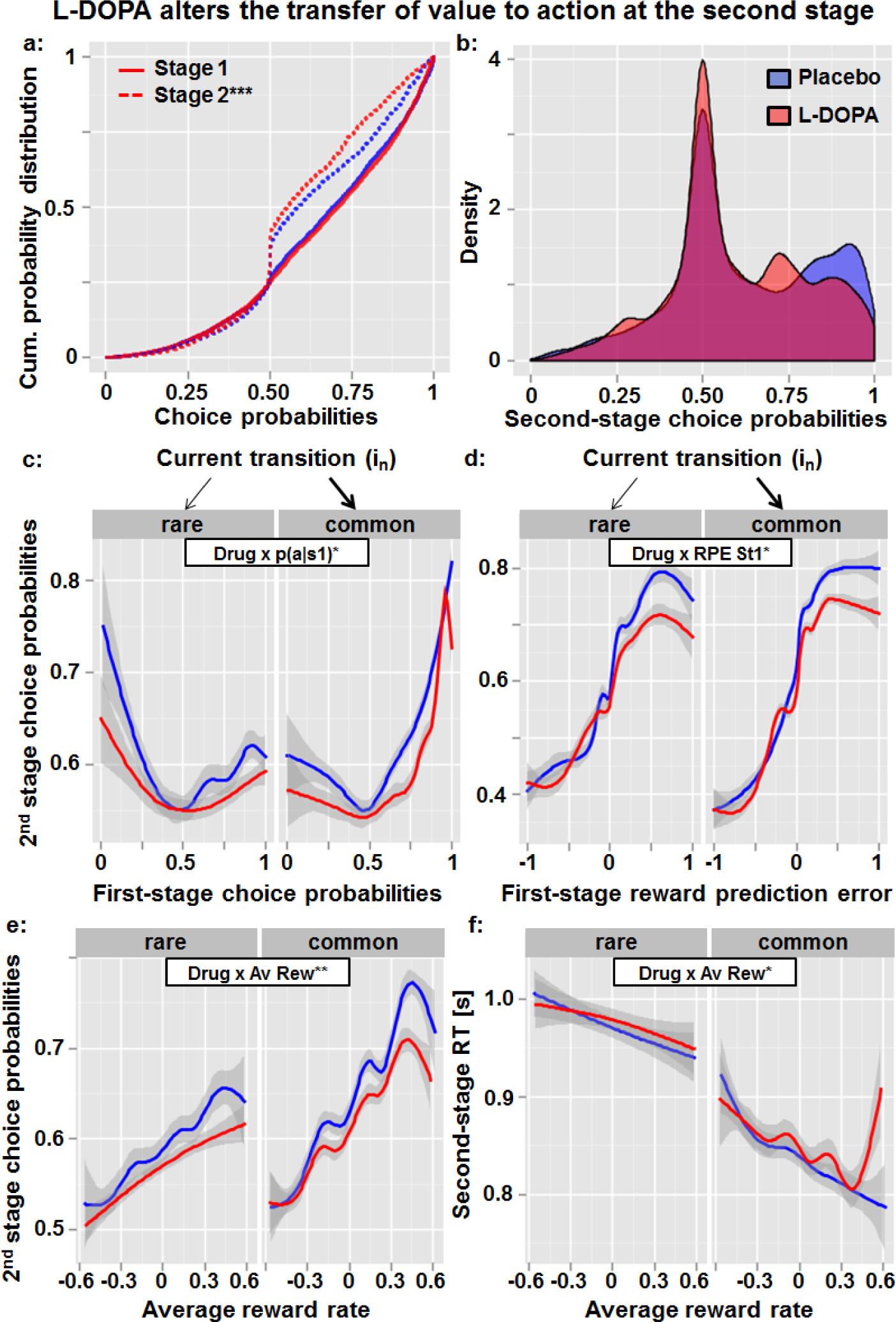
A: L-DOPA reduces stochasticity of choices at the second (i.e., model free*;p* < .001), but not the first stage (*p* = .52). B: Group density plots indicate that the net effect is mainly due to a reduction of choices with very high probabilities given the model’s estimation of choice values. C: The reduced correspondence of first-stage choice probabilities (*p* = .014) and (d) reward prediction errors with second-stage choice probabilities (*p* = .019) is not affected by transition and apparent for both rare and common transitions (Drug x Transition *p* > .27). E: The average reward rate (Av Rew), which reflects the recent history of wins (centered on each individual’s average percentage of wins), has less of an effect on second-stage choice probabilities after administration of L-DOPA (*p* = .003). F: The increase in response time at a high average reward rate in the L-DOPA condition (*p* = .048), particularly in common trials, suggests that instead of a reflexive invigoration of behavior, a recent stream of wins may have facilitated strategic exploration. This might add to the increased stochasticity of choices (noisy exploration) at the second stage. For the fitted models, see SI.

At stage 2, we observed a decrease in choices at the deterministic end (i.e., with very high probabilities; Figure 6a-b), which is reflected in the *β*2 parameter in the computational analysis. This observation was corroborated by the fact that L-DOPA reduced the correspondence between the estimated evaluations of options and resulting actions. The increased independence of action from value with higher dopamine tone was evidenced by the reduced correspondence between first-and second-stage choice probabilities (Figure 6c) and the reduced correspondence between first-stage prediction error signals and second-stage action values (Figure 6d).

Furthermore, we assessed how increases in dopamine tone affect action control by the average reward rate, which has been linked to endogenous fluctuations in dopamine tone before^1^. In line with the predictions of the thrift theory, L-DOPA reduced the correspondence between the average reward rate and choice probability at second stage (Figure 6e).

Moreover, L-DOPA led to a characteristic increase in second-stage RT when the average reward rate was high and, hence, uncertainty is expected to be low (Figure 6f). This increase was not seen during placebo sessions. Here, RT decreased monotonically with a recent stream of wins in the task as one would expect when past success is being exploited to maximize wins. Collectively, our data suggests that L-DOPA reduces MF control by facilitating “noisy” and “strategic” exploration during the task.

### L-DOPA does not change overall task performance

As we have detailed before, L-DOPA had no overall effect on the obtained model fit of the computational model of choice and on general RT. However, given that L-DOPA increases stochasticity of choices at the second stage of the task and facilitates exploration; does it come at the cost of the number of obtained rewards? We found that L-DOPA tended to increase the rate of rewards obtained during the task (*t* = 1.70, *p* = .093; M_D_ = 48.8% vs. M_P_ = 47.7%). Importantly, the rate of rewards was positively correlated with the fit of the computational model (*r*_D_ = .34, *p* = .006; *r*_P_ = .42, *p* = .001) indicating that the model reflects differences in task performance. Consequently, there was little indication that the observed reduction in MF control was disadvantageous in terms of overall task performance.

## Discussion

Tonic and phasic dopamine signaling are known to play key roles in reinforcement learning and action control. In the current study, we found that increases in dopamine tone induced by L-DOPA reduced the reflexive MF control of behavior via reduced direct reinforcement of successful actions. In contrast, deliberative MB control of behavior was unaffected by L-DOPA. The changes in MF control of behavior were not explained by changes in learning rates at different stages of the task or for positive versus negative outcomes. Moreover, these changes were not explained by differential coding of RPE signals as we did not observe differences in their correspondence with BOLD response. Nevertheless, in line with behavioral effects, L-DOPA reduced the effect of reward in the brain. First, it reduced the reward effect in the mesocorticolimbic system when the outcome (reward vs. no reward) was presented. Second, it reduced the facilitating effect of preceding reward on BOLD response in general at the beginning of the next trial. Collectively, these results conclusively indicate that L-DOPA reduces MF control of behavior by reducing the transfer of learned value to overt actions, which is in line with the recent value^16^ and thrift theories^5^ of dopamine function. This interpretation is corroborated by increases in noisy and strategic exploration, which is one of the unique predictions of the thrift theory^5-7^ that has not been extensively tested in humans before. Hence, our results add to the growing evidence for a dopaminergic contribution to exploratory decisions^7, 32^.

Our main behavioral finding that L-DOPA reduces MF control without affecting MB control of behavior is in line with the observation that behavioral effects of phasic dopamine signals are dependent on their tonic baseline^16, 27^, which prominently modulates the signal-to-noise ratio of phasic burst signaling^33, 34^. MF control is mainly driven by phasic firing of dopaminergic neurons encoding temporal difference RPEs^8, 9, 35^. We observed that RPE signals and learning rates were not altered by an increase in dopamine tone, which suggests that reward learning *per se* was not affected. Whereas Pessiglione, Seymour, Flandin, Dolan and Frith^19^ had reported increased amplitudes of RPE signals after L-DOPA that explained improvements in reward learning, these effects were only significant compared to the haloperidol group. Furthermore, in line with our results, Wunderlich, Smittenaar and Dolan^13^ did not observe significant effects of L-DOPA on learning rates. Consequently, an increased baseline of dopamine may reduce the throughput of phasic signals since the local change in dopamine is attenuated, which in turn might reduce their potential to reflexively adjust actions. Such a decoupling between the dopaminergic mechanisms of reward learning and action control with heightened tone has been conclusively demonstrated in animal models^16, 17^ When alterations in behavior occur without resultant changes in reward learning^16, 36^, the correspondence between action and learned value is effectively reduced^36^. Notably, Wunderlich, Smittenaar and Dolan^13^ also reported an (non-significant) increase in choice stochasticity for L-DOPA. Since they only reported drug effects for the reduced model that did not differentiate choice stochasticity between the two stages of the task, it is conceivable that this could have masked a significant effect of L-DOPA given the marked Stage x Drug interaction on stochasticity that we observed.

Notwithstanding, the absence of L-DOPA effects on MB control appear to be in contrast with the findings reported by Wunderlich, Smittenaar and Dolan^13^, or Sharp, Foerde, Daw and Shohamy^37^. However, several aspects may contribute to the observed discrepancies. First, when we included the obtained fit of the computational model during the L-DOPA session as a covariate, we found that MB control only increased in participants whose behavior was well approximated by the model. Likewise, when we entered working memory capacity as a covariate, we found that MB control only increased in participants with high working memory capacity. Thus, it is possible that our representative sample of adults performed the task less in line with the computational model or at a lower overall level, which could reduce the net effect of the drug in the studied sample. In other words, this suggests that facilitating effects of L-DOPA on MB control do not generalize to the population because they could be mediated by interindividual differences in working memory capacity^31^ or, perhaps, presynaptic dopamine levels in the ventral striatum^14^ that are known to contribute to overall task performance. If we were to restrict our analysis to individuals with high working memory capacity, we could essentially replicate that L-DOPA increases MB control^13^. Second, effects of L-DOPA on the MF vs. MB control weighting parameter ω were inconsistent across the alternative computational models, which may reduce reproducibility of results in general. Importantly, the reduction in MF control of behavior was observed independently of the fit and the parametrization of the computational model suggesting that this was a general effect of increased dopamine tone. Third, both previous samples were small (N_1_ = 18; N_2_ = 22) and given our effect-size estimate of reduced MF control (r ~.30), the power to detect such an effect would be below 30%. In line with this interpretation, both studies reported decreased MF control which was, however, not significant. Collectively, these results suggest a complex interaction between tonic and phasic dopamine signaling on the one hand and the arbitration between MF vs. MB control on the other hand. Whereas L-DOPA may reduce MF control in general yet to a moderate degree, the net effects on MB control are perhaps mediated by interindividual differences in task performance and working memory.

Critically, the behavioral effects of L-DOPA on MF control were also echoed in BOLD response. In line with reduced MF control of behavior, we observed an attenuation of the reward outcome signal in the mesocorticolimbic system and a global attenuation of the effect of preceding reward at the onset of the subsequent trial. Moreover, in the ACC, L-DOPA abolished the correspondence between reward outcome signals and MF control of behavior. Taken together, these findings suggest that L-DOPA reduces the transfer of value to action, which is well in line with previous studies. In patients with Parkinson’s disease, L-DOPA (on medication condition) disrupts performance of reversal learning^28^ and NAcc activation during reversal learning, particularly when the final reversal error occurs (i.e., preceding a switch^38^), which might be indicative of reduced transfer of value. Moreover, L-DOPA has been shown to reduce habit learning (i.e., MF control) in the weather prediction task in patients off medication^39^. Likewise, L-DOPA decreased the influence of a “Pavlovian controller” on instrumental action in healthy individuals such that action learning became less affected by outcome valence^40^ and it increased a value-independent propensity to gamble by making risky options more attractive to pursue^41^. Thus, increases in tonic dopamine may allow an individual to “escape” their previous reward history by embarking on a more effortful exploration of available options in the environment^5, 7^. It is plausible that this neurobiological mechanism might also account for reported increases in switching between tasks^28^ and this hypothesis calls for future research on a more comprehensive set of tasks.

The current study has limitations that will need to be addressed in future research. First, whereas it is commonly assumed that the dopamine precursor L-DOPA primarily raises dopamine levels in the striatum, it is conceivable that the effects L-DOPA are not solely attributable to the transformation into dopamine and subsequent postsynaptic activation of dopaminergic receptors^42^. Second, we observed that L-DOPA made decisions more independent of the current evaluation of options, which would be expected to impair task performance. In contrast, we found that L-DOPA tended to increase the number of wins. Thus, it remains to be determined if L-DOPA facilitates exploratory behavior to an extent that is even advantageous in environments characterized by volatile reinforcement schedules.

To conclude, we found that increases in dopamine tone lead to decreases in reflexive MF control of behavior by reward while deliberative MB control, which takes the learned transition structure of the environment into account, and value-based learning remained unaffected on average. These results suggest that L-DOPA reduces the transfer of learned value to action, which may result from the reduced local change in dopamine induced by phasic dopamine release when tonic levels are heightened. Our observations corroborate recent value^16^ and thrift^5^ theories of dopamine function pointing to an essential role of dopamine in supporting the invigoration of responses and energy expenditure^43^. As a result, heightened dopamine tone may allow an individual to break reflexive and habit-like chains of action, which have been established because of their previous reward history, and support exploration of available options in the environment^5, 7^. Hence, behavioral flexibility may arise as a consequence of increased independence of future behavior on the preceding stream of success.

## Methods

### Participants

This dataset is part of an ongoing study investigating dopaminergic modulation of reinforcement learning. In order to ensure that the participants of our study were representative of the general population, we requested for postal addresses of individuals randomly selected by the residents’ registration office of Dresden, Germany (N = 15,778) and invited them to our study. To limit a confounding effect of age^44^, we recruited participants within the age range of 30-40 years. As part of the general study protocol, participants were invited to a total of four visits, which comprised of a pre-screening visit and two fMRI visits at the Neuroimaging Center, Technische Universität Dresden, followed by a positron emission tomography (PET) visit at the PET center (data reported elsewhere). For the current analysis, we included 65 healthy participants (49 male; *M*_*age*_ = 37.0 years, *SD*_*age*_ ± 3.56, range [30-42]) who completed both drug and placebo sessions without severe side effects (N = 10 had only one visit; *N* = 5 reported severe side effects in the second session) and passed extensive quality control (N = 4 were excluded because of low fMRI data quality, *N* = 2 because of >20% missing trials in one session). Furthermore, four participants had to be excluded due to 1) brain atrophy, 2) a positive drug test for THC, 3) data loss, 4) erroneous drug manipulation. This sample size provides sufficient power (1 – β = .80) to detect small- to medium-sized effects of repeated measures at the behavioral and ROI level (α = .05, dz = .36) and medium-sized effects at the voxel level (α = .001, dz = .54).

Inclusion criteria for the study were as follows: (1) at least 30 years old at the date of the PET scan, (2) no history of neurological or mental disorders according to the Screening Version of the Structured Clinical Interview for DSM-IV (Wittchen et al., 1997) except for nicotine dependence, (3) no MRI, PET nor L-DOPA contraindications, (4) normal or corrected-to-normal vision, (5) no recent use of illicit drugs (urine test on first fMRI visit; Kombi/DOA10-Schnelltest, MAHSAN Diagnostika GmbH, Reinbek, Germany) nor alcohol consumption (breath-alcohol analysis on both fMRI visits; Alcotest 6510, Drägerwerk AG & Co. KGaA, Lübeck, Germany). Since the main goal of recruitment was to maximize generalization to the population, we deliberately included smokers (~30% of the adult population in Germany). Smokers were allowed to smoke cigarettes prior to the fMRI visit and breath carbon monoxide levels were measured to assess recent use of cigarettes. The institutional review boards of Technische Universität Dresden approved the study and we obtained informed consent was obtained from all participants prior to taking part in the experiment.

### Paradigm

Participants performed an adapted version^45^ of the two-stage Markov decision task developed by Daw et al. while undergoing fMRI. For the current study, (a) the instructions were translated into German, (b) visual stimuli were adapted to present different sets of stimuli across visits (pseudo-randomized across participants), and (c) outcome presentation times at both stages were decreased by a factor of 2 to reduce trial duration. The task consisted of a total of 201 trials, separated by inter-trial intervals sampled from an exponential distribution (M = 2 s; range: 1-7 s). For each trial, there were two stages (Figure 1a). At the first stage, participants had to choose between two grey boxes. After the first-stage choice, they were led to a second stage where they had to make a choice between two colored (green/yellow) boxes. After they have made the second-stage choice, the monetary outcome (win 20 cents or 0 cents) for the trial was presented. In order to optimize their performance, participants had to learn two aspects of the task. First, they had to learn the transition structure, that is, which grey stimulus led to the yellow pair of stimuli in 70% of the trials (“common” trials) and to the green pair in 30% of the trials (“rare” trials; and vice versa for the other grey stimulus). Second, they had to infer the reward probabilities associated with each second-stage stimulus, which followed random Gaussian walks that changed slowly and independently of each other with reflecting boundaries at 0.25 and 0.75. Both aspects were emphasized in the instructions and participants completed 50 practice trials using an independent set of stimuli. After participants completed the practice, they were queried to ensure that they understood a) the transition structure and the difference between common and rare transitions and b) that the best option changes over time due to the random walks. If participants did not answer the query questions correctly, the experimenter repeated the instructions.

This task has been employed previously in multiple studies for characterizing weighted contributions of MF and MB systems in individuals during adaptive learning e.g.^3,13,14, 45, 46^. The key feature differentiating between MF and MB strategies is how first-stage choices are influenced by the “model”, that is, the transition structure between the two stages of the task. For example, suppose an individual was rewarded for a second-stage choice during a rare transition trial. In order to be rewarded again for the same second-stage option, MF individuals would repeat their first-stage choice simply because it was rewarded. On the other hand, MB individuals would consider the transition probabilities of both first-stage stimuli. Hence, they would switch to the *other* first-stage stimulus. The reason is that according to the transition structure, switching to the other first-stage option would give them a higher chance to select the same second-stage stimulus, which increases the chances of being rewarded again^3^.

### Procedure

During the intake visit, we measured height and weight, drew blood samples, and participants completed tasks and questionnaires at a computer. Participants then returned to the scanning facility for their fMRI visits. In order to minimize the influence of medication on BOLD signal, participants were asked to abstain from medication for at least 24 h prior to their visit. As presence of food in the bowels influences the rate of levodopa absorption^47^, we wanted to control for the amount of food present by asking participants to fast overnight before arriving at the scanning facility. Participants were then given a small standardized breakfast (about 25 g butter biscuits, ~120 kcal) upon arrival (between 05:30 – 08:45 am) and dextrose tablets throughout the session (about 4 g, ~17.4 kcal/h) to reduce side effect of the drug administration.

Next, participants were instructed and trained on the task. In line with previous studies, we explained that (a) transition contingencies (mapping between first and second stage) would remain fixed throughout the experiment, (b) reward probabilities of each option (“states”) at second stage would vary slowly over time independently of each other, and (c) they should try to maximize their monetary outcome throughout the experiment. To familiarize the participants with the task, they were given a computerized practice that consisted of 50 trials prior to the scanning session with the same transition contingencies (70%/30%). To minimize transfer of expectations to the experiment, the practice task had a different set of reward probabilities and stimuli from that used inside the scanner.

Furthermore, participants were asked to complete a series of computerized and pen-and-paper questionnaires on psychological functioning (e.g., mood). On the second fMRI visit, they completed working memory tests instead of being trained on the two-stage Markov task. After approximately 80 min, we administered 150mg/37.5mg L-DOPA/benserazide (Madopar; Levodopa and Berazidhydrochlorid; Winthrop Arzneimittel) orally following a double-blind, placebo-controlled (P-Tabletten, Lichtenstein; Roche) randomized cross-over design. The order of drug condition during the fMRI visits was pseudo-randomized across participants prior to the study. Participants then entered the scanner for structural scans and a second blood sampling (T1) before they proceeded with the two-stage Markov task, which took about 36 min.

After participants completed the two-stage Markov task, a 6-min resting-state scan was collected. The resting-state scan was followed by a booster dose of L-DOPA, a second task inside the scanner, and additional behavioral testing outside the scanner. Participants took approximately five hours to complete one session. At least seven days after the first fMRI visit (M = 12.2 d, ±9.6 d), participants returned and completed a second visit following the same procedure (except for exchanging the task training for a set of working memory tests), but receiving the complementing drug condition instead.

### fMRI data acquisition and preprocessing

MRI images were acquired on a 3 Tesla Magnetom Trio Tim system (Siemens, Erlangen, Germany) equipped with a 32-channel head coil. During the in-scanner task, stimuli were presented on an MR compatible screen and rearview mirror system. Participants responded by pressing their index fingers on two separate button boxes, one held in each hand. Psychophysics Toolbox Version 3^48, 49^ implemented within MATLAB R2010a software (The Mathworks, Inc., MA, USA) was used to present the stimuli and collect behavioral data. Functional images were acquired using a gradient echo-planar imaging (EPI) sequence, repetition time TR = 2.41 s; echo time TE = 25 ms; flip angle: 80°; field of view: 192 x 192 mm^3^; matrix size: 64 x 64; voxel size: 3 x 3 x 2 mm^2^ (slice thickness: 2 mm; gap: 1 mm). Every volume consisted of 42 transverse slices acquired descending from the top, manually adjusted ~25° clockwise from the anterior commissure-posterior commissure plane (total ~900 volumes for each participant, total scan time: ~36min). A corresponding field map was also recorded for distortion correction of the EPI images. Structural images were acquired using a T1-weighted magnetization prepared rapid acquisition with gradient echo (MPRAGE) sequence for normalization, anatomical localization as well as screening for structural abnormalities by a neuro-radiologist (TR: 1.90 s; TE: 2.52 ms; flip angle: 9°; field of view: 256 x 256 mm^2^; number of volumes: 192; voxel size: 1 x 1 x 1 mm^3^).

Functional brain data were preprocessed using SPM8 (Wellcome Trust Centre for Neuroimaging, London, UK) implemented within Nipype Version 0.9.2^50^. The first 4 volumes of the EPI images were discarded to allow for magnetic saturation. The remaining 896 volumes were subjected to slice-time correction (reference: middle slice), followed by realignment to the first volume of the run to correct for motion. Distortion correction based on the field map was then applied to the realigned EPI images. Each individual anatomical T1 image was first co-registered to the individual mean EPI image before segmentation and normalization to MNI space. The resulting transformation parameters were then applied to the distortion-corrected EPI images to spatially normalize them to MNI space (non-linear; resampled to 2 x 2 x 2 mm^3^). Finally, normalized EPI images were spatially smoothed with an isotropic Gaussian kernel (full width at half maximum = 8 mm). During first-level analyses, the data was high-pass filtered at 128 s. As mentioned previously, four participants were excluded from the final analysis because of low fMRI data quality due to excessive in-scanner motion (N = 2, > 3 mm translation or 3° rotation volume-to-volume), anatomical abnormality (N = 1), and failure in image segmentation (N = 1).

### Data analysis

#### Factorial analysis of model-free versus model-based behavior and response time

We investigated the effects of increases in tonic dopamine on stay/switch behavior using the task conditions (reward, transition) in a factorial analysis of the two-stage Markov decision-task (Figure 1)^3, 13, 14^. We had two behavioral measures of interest, namely the tendency to select the same choice as in the previous trial (“stay”) and response time (RT). For this set of behavioral analyses, we estimated the main effects of reward, transition, and the Reward x Transition (RxT) interaction on the repetition of the same choice at the first stage of the next trial using full mixed-effect logistic regression analysis of placebo and drug sessions as implemented in hierarchical generalized linear modeling (HGLM, outcome distribution Bernoulli). The two main effects and the interaction term were treated as random effects, that is, we computed the deviation of each individual from the group effect for drug and placebo sessions to freely estimate individual values of the effects. To test for significant differences, we used empirical Bayes (EB) estimates that take group priors into account and ordinary least squares (OLS) estimates, which are not affected by group priors. We used both estimates because OLS values are “unbiased” and provide a lower-bound estimate of the group effect, but they come at the cost of power. This is because the estimates are not shrunk based on the likelihood of observed values across the group, unlike empirical Bayes estimates, leading to deviations from the normal distribution. For the second set of behavioral analyses, we set up two separate models for RT at first stage and RT at second stage. To estimate factorial effects on RT, we log transformed RT to normalize the distribution and used hierarchical linear modeling (HLM). The RT model at first stage was set up analogously to the choice model. The RT model at second stage included first-stage RT and the current transition as additional predictors. Both RT models included trial number as a random effect to account for potential reductions in RT across the run. All these analyses were conducted with HLM 7^51, 52^.

#### Computational model

As detailed by Daw, Gershman, Seymour, Dayan and Dolan^3^, we assumed that agents learn by updating state-action values at each trial, *t*, through a weighted combination of MB and MF components. The MF component learns the system using a temporal difference algorithm, whereas the MB component does so by maximizing the expected value by taking the contingencies into account.

The second stage, *S*^*’*^, has only an MF component driven by the final reward, *r*_*t*_,

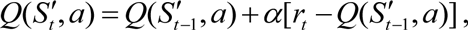

where 0 ≤ *α* ≤ 1 is the learning rate. The MF component of the state-action values at the first stage, S, is updated similarly with the addition of an eligibility trace parameter, 0 ≤ *λ* ≤ 1. However, the presence of different transition probabilities (the task “model”) leads to a potential MB learning component as

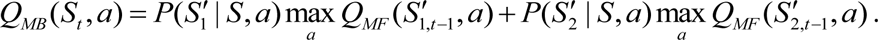

The overall state-action values at the first stage are computed as a linear combination of the MF and MB components,

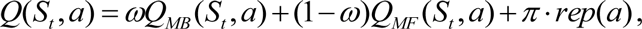

in which 0 ≤ *ω* ≤ 1. The tendency to stick to the first-stage action taken at the previous trial is captured by a perseveration parameter *π*. In order to choose an action, a softmax function maps the state-action values to choice probabilities at every stage as

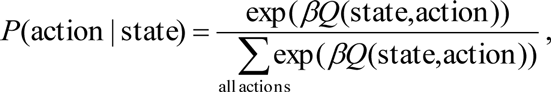

in which *β* ≥ 0 is the so called inverse temperature. Thereby, it represents choice stochasticity, that is, how strongly choices made are related to expected values. The strength of choice consistency is related to the exploration/exploitation trade-off^53^ and higher values correspond to stronger exploitation of learned values. Considering different learning rates and stochasticity parameters for the two stages, the model comprises seven parameters.

Other variants of reinforcement learning models have been studied for this task before and further details are provided in the SI. Wunderlich, Smittenaar and Dolan^13^ investigated the effects of simplifying the original model to have four parameters (*α*, *β, ω, π)* and, alternatively, considering different learning rates for positive and negative prediction errors (*α*_+_, *α*_−_, *β*, *ω*, *π*). Recently, new formulations have been presented, which decouple MB and MF components and relax the assumption of a linear weighting^30, 31^ . Furthermore, an alternative is provided by integrated reinforcement learning architectures such as DYNA, which assume that behavior is completely controlled by the model-free system. In this model model-based system only has an indirect influence by training the model-free system offline using simulations of the state space of the task^54^ However, due to our a priori hypotheses, which were focused on the original formulation of the model, we did not evaluate these alternatives in detail in the current study.

Finally, for any individual, we sought to find a set of parameters which yields the highest likelihood of the data for a given model. Here, we used maximum a posteriori estimation (*MAP)* to improve the estimation of parameters for parametric statistics^14^.

#### Statistical modeling of fMRI data at first level

First and second-level analyses of fMRI data were carried out using SPM8. For this study, we calculated two sets of first-level statistics that correspond with each of the behavioral analyses. The first set of analyses was conducted in accordance to Daw, Gershman, Seymour, Dayan and Dolan^3^. The regressors are based on the computational model of behavior e.g.,^14, 46^ and use the mean parameter values of the computational model across both sessions. Briefly, it incorporated the onset of stage 1 as event with the parametric regressors first-stage action values, *P*(a_1_,_t_|s_A_), and its partial derivative with respect to ω, a combined event regressor for the onset of stage 2 and the onset of outcome presentation with the parametric regressors MF RPEs and MB RPEs. The two events were merged to enable the conjoint estimation of MF and MB RPEs on one onset regressor. To capture potential differences in BOLD response between the two events, the onset of the outcome event was modeled separately in addition. Parametric predictions of MF and MB RPE were derived for both event onsets given the assumption of absolute MF (ω = 0) or MB (ω = 1) control, respectively. Notably, the MB regressor was set up to capture only variance that is not accounted for by MF RPEs since MB RPEs are assumed to act on top of MF control^3^. As there is no transition involved at the second stage, reinforcement learning is primarily under MF control when the outcome is being presented and MB RPEs are therefore set to 0.

For the second set of first-level statistics, we employed a factorial approach that mirrors the behavioral analysis of stay/switch behavior. Here, we used a regressor for the onset of stage 1 with the parametric regressors previous reward (coded as -0.5 / 0.5), previous transition (coded as -0.5 / 0.5), and their interaction, a regressor for the onset of a common transition trial at stage 2, a regressor for the onset of a rare transition trial at stage 2, a regressor for a reward at outcome onset (i.e., the result of the choice at stage 2), and a regressor for reward omission at outcome onset. Specifically, we used the contrast rewarded – unrewarded trials at outcome onset and the parametric effect of preceding reward at first-stage onset to evaluate the simple main effects of L-DOPA on processing of reward, which were tested based on the observed behavioral differences between drug conditions.

For group inferences, we computed second-level group statistics with placebo vs. L-DOPA as repeated measures factor for the parametric contrast images (MF RPE, MB RPE, first-stage action values, rewarded vs. unrewarded trials). The second-level statistics based on the computational model included order as a nuisance variable. Since the effects of order were negligible, we included only the estimated effect of the reflexive effect of reward on stay behavior when we evaluated brain-behavior interactions.

ROI analyses were focused a priori on the NAcc and vmPFC where both types of signal correlates are evident^3,14^. Valence effects of reward and MF RPE signals were also expected a priori to occur in the VTA/SN (e.g., see the term “reward” at www.neurosynth.org;^55^). For brain–behavior coupling, we furthermore included the ACC since it is critically involved in the allocation of effort according to learned action policies^56, 57^

#### Full mixed-effects modeling of time courses

To maximize the sensitivity in detecting potential drug effect on RPE signals, we complemented the common so-called summary statistic random effects analysis by implementing a full mixed-effects design (i.e., two-level HLM). Here, we estimated drug effects by incorporating group priors based on extracted ROI time series as described by Kroemer, Guevara, Ciocanea Teodorescu, Wuttig, Kobiella and Smolka^58^. To this end, we extracted the first eigenvariates from anatomical masks of the nucleus accumbens (NAcc) and ventromedial prefrontal cortex (vmPFC) where both MF and MB RPE signals can be observed. This method improves sensitivity of parametric statistics similar to MAP estimation for the behavioral data. To estimate drug effects on variables of interest, we allowed for interactions of L-DOPA with MF RPEs, MB RPEs, and first-stage action values. All events, parametric regressors, and interactions of L-DOPA with parametric regressors were modeled as random effects at the participant level and, thereby, led to parameter estimates for each individual.

#### Full mixed-effects modeling of exploratory behavior (as depicted in Figure 6)

We evaluated choice probabilities and RT in more detail using the fitted predictions of the computational model (M_7P_) obtained after the MAP estimation for each individual participant. We concatenated data from placebo and drug sessions and predicted choice probabilities at the first (Figure 6a-b) and second stage (Figure 6a-e). For Figure 6f, we predicted second stage RT as the outcome (ln transformed to meet distributional assumptions). The setup and output of the full-mixed effects models is detailed in the SI (Table S.4). The average reward rate was computed by using a Gaussian moving window incorporating the past 5 wins according to a recency-weighting scheme and varied between 0 (5 omissions in a row) and 1 (5 wins in a row). For statistical analyses, we centered the reward rate for each individual session. Thus, the regressor captures residual variance throughout a session that is accounted for by the recent stream of success.

#### Statistical threshold and software

For behavioral and ROI analyses, we used α = .05 (two-tailed) as significance threshold and performed correction for multiple comparisons based on the hypothesis test as detailed in the results section. For fMRI whole-brain analyses, we used one-sided contrast maps (mass-univariate *t*-tests) thresholded at *p* < .001 to assess cluster size. To facilitate the visualization of the involved brain structures, we show the images at a slightly lower threshold as indicated in the figures. To check for robustness, key parametric results were also assessed with non-parametric equivalents and corresponding p-values were derived by Monte Carlo simulations. To analyze and plot data, we used SPSS v21-23, R v3.2.2^59^, R Deducer^60^, HLM v7^51, 52^, Mango v3.6-3.8, and MATLAB v2012-2015.

## Acknowledgement

We thank Annika Kienast, Elisabeth Kiese, and Valerie Fournes for help with data acquisition as well as Michael Marxen and Dirk Müller for support in preprocessing fMRI data. Furthermore, we thank Daniel Schad for insightful discussions regarding data analysis. The study was supported by the Deutsche Forschungsgemeinschaft, grants SFB 940/1, 940/2

## Author contributions

MNS and TG were responsible for the study concept and design. YL collected data.

SP set up the computational modeling. NBK performed the data analysis and YL and SP contributed to analyses. NBK wrote the manuscript. All authors contributed to the interpretation of findings, provided critical revision of the manuscript for important intellectual content and approved the final version for publication.

## Financial disclosure

The authors declare no competing financial interests.

